# *Ccdc38* is required for sperm flagellum biogenesis and male fertility in mouse

**DOI:** 10.1101/2022.01.10.475757

**Authors:** Ruidan Zhang, Bingbing Wu, Chao Liu, Xiuge Wang, Liying Wang, Sai Xiao, Yinghong Chen, Huafang Wei, Fei Gao, Li Yuan, Wei Li

## Abstract

Sperm flagellum is essential for male fertility, defects in flagellum biogenesis are associated with male infertility. Deficiency of CCDC42 is associated with malformation of the mouse sperm flagella. Here, we find that the testis-specific expressed protein CCDC38 (coiled-coil domain containing 38) interacts with CCDC42 and localizes on manchette and sperm tail during spermiogenesis. Inactivation of CCDC38 in male mice results in distorted manchette, multiple morphological abnormalities of the flagella (MMAF) of spermatozoa, and eventually male sterility. Furthermore, we find that CCDC38 interacts with intra-flagellar transport protein 88 (IFT88) as well as the outer dense fibrous 2 (ODF2), and its depletion reduces the transportation of ODF2 to flagellum. Altogether, our results uncover the essential role of CCDC38 during sperm flagellum biogenesis, and suggesting the defects of these genes might be associated with male infertility in human being.

**Summary statement:** We demonstrated that CCDC38, localizes on manchette and sperm tail, is crucial for male fertility.

## Introduction

Sperm flagellum is essential for sperm motility (Freitas et al., 2017, Pereira et al., 2017), which is a fundamental requirement for male fertility. The flagellum contains four parts: connecting piece, midpiece, principal piece and end piece. The core of sperm flagellum is the central axoneme, which consists of a central microtubule pair (CP) connected to 9 peripheral outer microtubule doublets (MD) to form ‘9+2’ structure (Sironen et al., 2020). The axoneme possesses radial spokes that connect the central and peripheral microtubules and are related to the mechanical movement of the flagellum (Inaba, 2011). Besides the axoneme, sperm flagellum contains unique structures, outer dense fiber (ODF) and fibrous sheath (FS) that are not present in cilia or unicellular flagella (Fawcett, 1975). The outer dense fibers (ODFs) are the main component cytoskeletal elements of sperm flagellum, which is required for the sperm motility (Inaba, 2011). ODFs contain 9 fibers in the midpiece, each of which is associated with a microtubule doublet. In the principal piece, ODFs 3 and 8 are replaced by two longitudinal columns of fibrous sheath (FS), in human, the diminished 3 and 8 fibers are finished at the annulus (Azizi and Ghafouri-Fard, 2017, Kim et al., 1999). There are at least 14 polypeptides of ODFs such as ODF1, ODF2 (Lehti and Sironen, 2017). Any defects in the axoneme structure can cause abnormalities in the sperm flagellum, change its morphology, causing severe sperm motility disorders (Sha et al., 2014). Thus, axoneme structures are very important to sperm morphology and the function of flagellum.

Multiple morphological abnormalities of the flagella (MMAF) is a kind of severe teratozoospermia (Coutton et al., 2015), which is characterized by various spermatozoa phenotype with absent, short, coiled, irregular flagellum and others. There are many flagellar axoneme defects in MMAF patients, including disorganization of microtubule doublets (MD), outer dense fibrous (ODF), fibrous sheath (FS), outer or inner dynein arms (ODA, IDA) and others (Jiao et al., 2021). Over the past several years, many mutations have been found to be associated with MMAF patients, and a lot of mouse models display MMAF-like phenotype, *Dnah2* (Li et al., 2019), *Dnah8* (Liu et al., 2020), *Cfap44, Cfap65* (Tang et al., 2017, Li et al., 2020), *Qrich2* (Shen et al., 2019), *Cep135* (Sha et al., 2017) and *Ttc21a* (Liu et al., 2019) are reported to MMAF related genes. Despite rapid progress in understanding the mechanism of MMAF, the pathogenesis of many idiopathic MMAF patients is still unknown.

The coiled-coil domain-containing (CCDC) proteins are involved in a variety of physiological and pathological processes. Increasing number of CCDC proteins have been suggested to be involved in ciliogenesis (Priyanka and Yenugu, 2021). But only some of those genes are involved in spermatogenesis, such as *Ccdc9, Ccdc11, Ccdc33, Ccdc42, Ccdc63, Ccdc172*, which are associated with sperm flagellum biogenesis and manchette formation, their defects lead to male infertility (Sha et al., 2019, Wu et al., 2021, Tapia Contreras and Hoyer-Fender, 2019, Young et al., 2015, Yamaguchi et al., 2014). *Ccdc42* is highly expressed in mouse testis, it localizes on manchette, HTCA and sperm tail during spermatogenesis, and it is necessary for HTCA assembly and sperm flagellum biogenesis (Tapia Contreras and Hoyer-Fender, 2019). However, the functional role of CCDC42 in spermatogenesis is still poorly understood.

Here, we found that CCDC38 was directly interacted with CCDC42, and it was expressed in the testis, and associated with the manchette in elongating spermatid. Importantly, *Ccdc38* knockout in mice resulted in abnormally elongated manchette and MMAF-like phenotype. Furthermore, we found that CCDC38 could interact with IFT88 and ODF2 to facilitate ODF2 transportation in flagella. Our results suggested that CCDC42 incorporating with CCDC38 mediates ODF2 transportation during flagellum biogenesis, and both are essential for flagellum biogenesis and male fertility in mice, suggesting some mutations of these two genes might be associated with male infertility in human being.

## Results

### CCDC38 interacts with CCDC42

Many CCDC proteins participate in flagellum biogenesis during spermiogenesis (Priyanka and Yenugu, 2021). CCDC42 localized to the centrosome, HTCA, manchette and sperm tail in male germ cells, and it is involved in the biogenesis of motile cilia and flagellum in mice (Perles et al., 2012, Tapia Contreras and Hoyer-Fender, 2019, Pasek et al., 2016, Silva et al., 2016). To understand the underlying mechanism of CCDC42 in flagellum biogenesis during spermiogenesis, we used STRING database to search for CCDC42-binding candidates (Fig. 1A). CCDC38, reported as a testis-specific protein (Lin et al., 2016), was chosen first. Epitope-tagged CCDC42 and CCDC38 expressed in HEK293T cells followed by immunoprecipitation experiments demonstrated that CCDC38 was detected in anti-MYC immunoprecipitates from CCDC42 co-transfectants, but not from cells co-transfected with the control plasmid (Fig. 1B). An overlapping immunostaining pattern was clearly found in Hela cells transiently expressing GFP-CCDC38 and MYC-CCDC42, and GFP-CCDC38 could also co-localized with γ-TUBULIN as reported (Firat-Karalar et al., 2014) (Fig. 1C). These results suggest that CCDC42 indeed could interact with CCDC38.

**Fig. 1.**
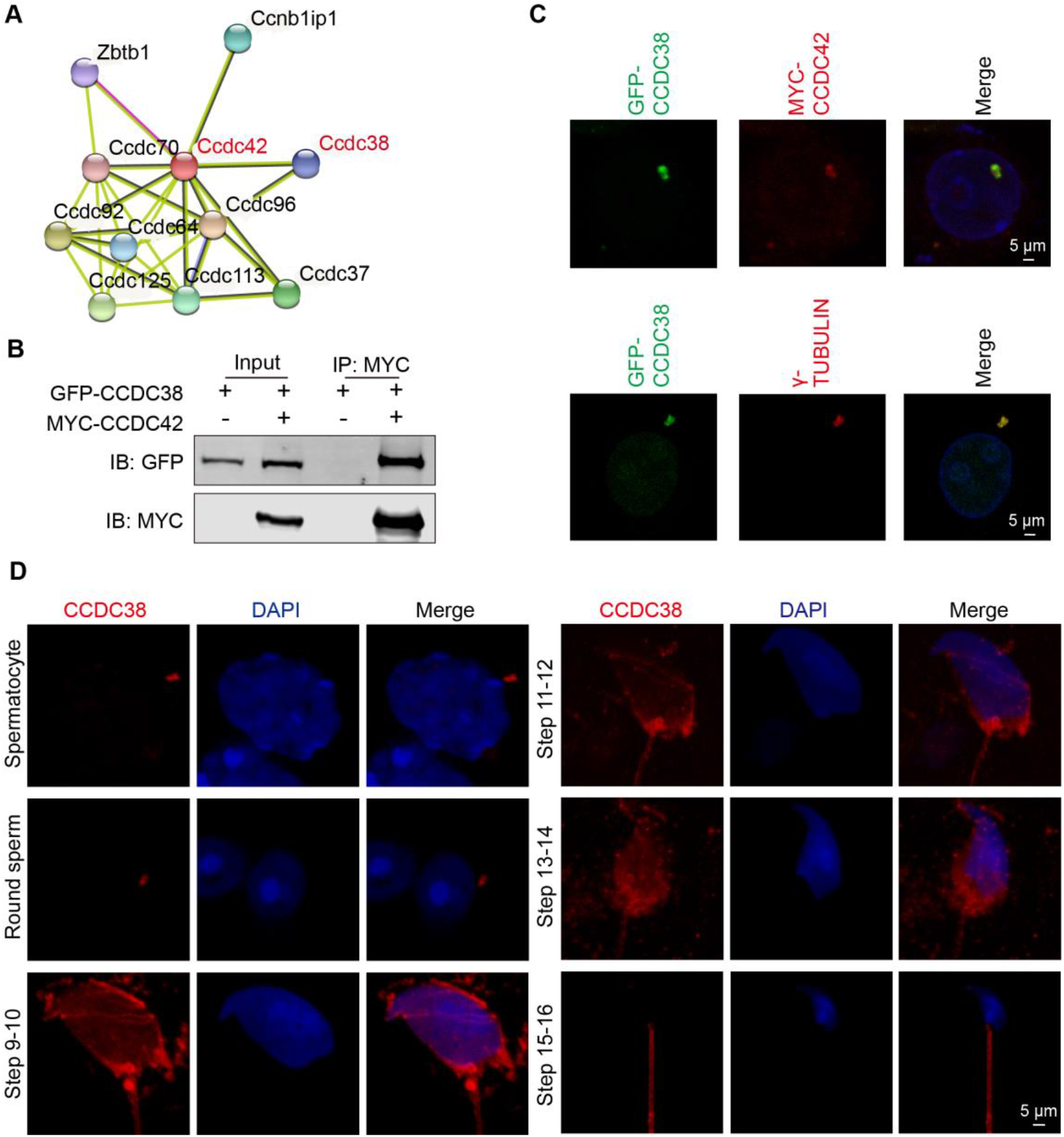
CCDC38 interacts with CCDC42. (A) CCDC38 might be interacted with CCDC42 predicted by the STRING database. (B) CCDC38 interacted with CCDC42. pCSII-MYC-CCDC42 were transfected into HEK293T cells with pEGFP-C1-CCDC38, forty-eight hours after transfection, cells were collected for immunoprecipitation with anti-MYC, and detected by anti-GFP or anti-MYC antibodies, respectively. (C) CCDC38 co-localized with CCDC42 and γ-TUBULIN in Hela cells. pCSII-MYC-CCDC42 and pEGFP-C1-CCDC38 were co-transfected into Hela cells. 48 h after transfection, cells were fixed and stained with anti-MYC and γ-TUBULIN antibody, and the nucleus was stained with DAPI. (D) The immunofluorescence of CCDC38 in WT mice. Testis germ cells were stained with anti-CCDC38 antibody, and nucleus was stained with DAPI.

Next, we examined the localization of the endogenous CCDC38 during spermatogenesis. CCDC38 was detected as two adjacent spots near the nuclei of spermatocytes or round spermatids, while it localized to the skirt-like structure encircling the spermatid head from step 9 to step 14 and the testicular sperm tail (Fig. 1D). We therefore speculate that CCDC38 might participate in flagellum biogenesis during spermiogenesis.

### *Ccdc38* knockout leads to male infertility

Reverse transcription-polymerase chain reaction (RT-PCR) revealed that *Ccdc38* was detected in the testis and firstly expressed at postnatal day 14 (P14), and peaked on P35 (Fig. 2A, B). To determine the physiological role of CCDC38, we generated *Ccdc38*-deficient mice by applying the CRISPR-Cas9 system to delete Exon 5 to Exon 11 of the *Ccdc38* gene (Fig. 2C). The *Ccdc38* knockout mice were genotyped by genomic DNA sequencing and further confirmed by PCR with 591 bp in *Ccdc38*^*+/+*^, and 750 bp in *Ccdc38*^*-/-*^ mice (Fig. 2D). Subsequent Western blotting analysis validated complete ablation of CCDC38 protein extracted from *Ccdc38*^*-/-*^ testes (Fig. 2E). We then examined the fertility of *Ccdc38*^*-/-*^ mice. Male *Ccdc38*^*-/-*^ mice exhibited normal mounting behaviors and produced coital plugs, but failed to produce any offspring after mating with WT adult female mice, in contrast, female *Ccdc38*^*-/-*^ mice generated offspring after mating with WT adult males (Fig. 2F). Surprisingly, the knockout of *Ccdc38* did not affect either testis size (Fig 2G) or the ratio of testis weight and body weight (Fig. 2H, I, J). Taken together, *Ccdc38* knockout leads to male infertility.

**Fig. 2.**
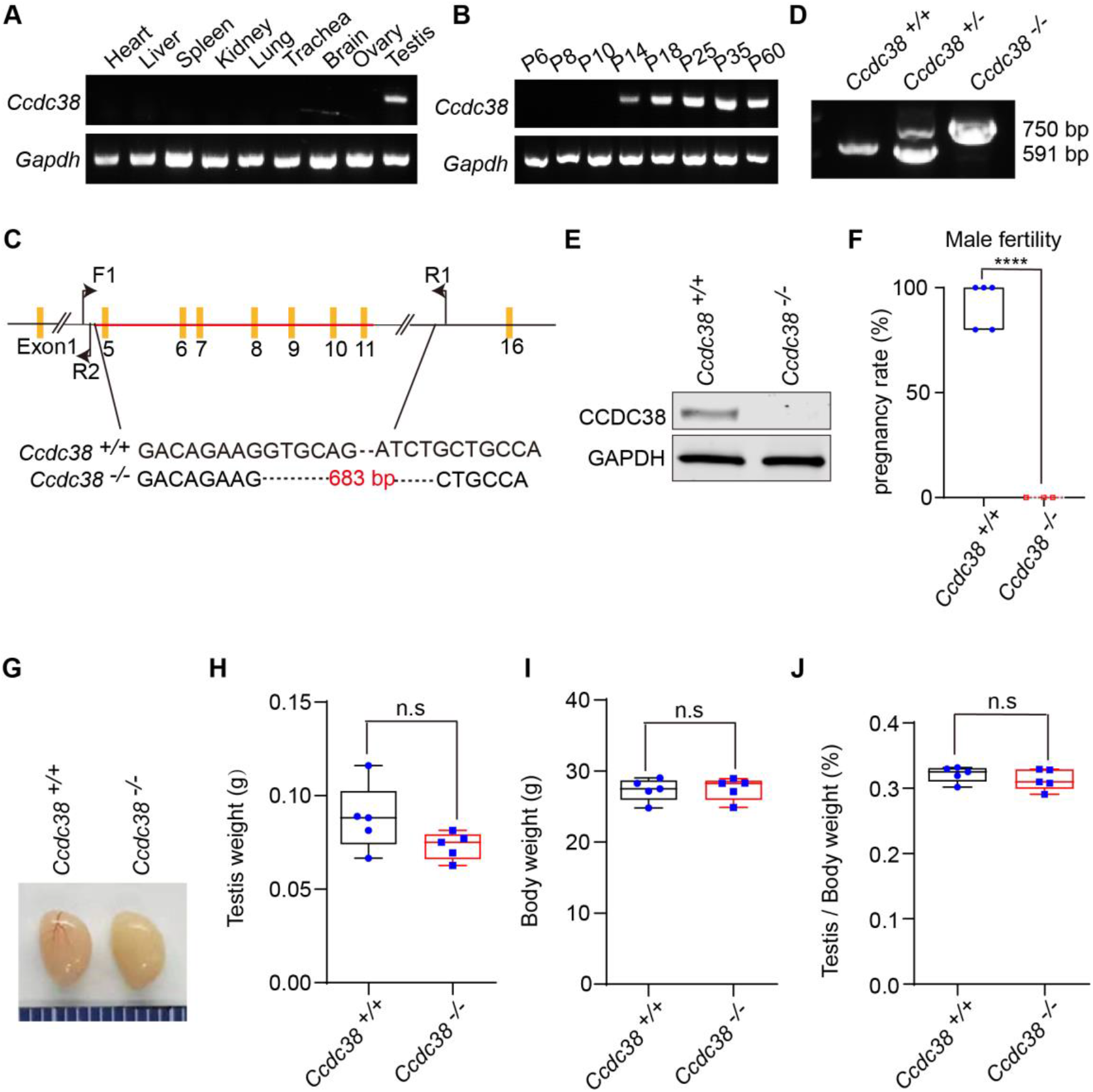
*Ccdc38* knockout leads to male infertility. (A) The expression of *Ccdc38* in different tissue. (B) The expression of *Ccdc38* in different days. (C) The generation of *Ccdc38*^*-/-*^ mice lacking exon 5-11. (D) Genotyping of *Ccdc38*^*-/-*^ mice. (E) Western blotting of CCDC38 indicated that the depletion efficiency in *Ccdc38*^*-/-*^ male mice. (F) The pregnancy rate of *Ccdc38*^*+/+*^ and *Ccdc38*^*-/-*^ mice at 2 month, there were no pregnancy mice in *Ccdc38*^*-/-*^ male mice. (G) The size of the *Ccdc38*^*+/+*^ and *Ccdc38*^*-/-*^ mice testes were not affected. (H) The testis weight in *Ccdc38*^*+/+*^ and *Ccdc38*^*-/-*^ male mice had no obvious difference (n=5). Data are presented as the mean ± SD. (I) The body weight in *Ccdc38*^*+/+*^ and *Ccdc38*^*-/-*^ male mice had no obvious difference (n=5). Data are presented as the mean ± SD. (J) The ratios of testis/body weight in *Ccdc38*^*+/+*^ and *Ccdc38*^*-/-*^ male mice were not affected (n=5). Data are presented as the mean ± SD.

### *Ccdc38* knockout results in MMAF

To further explore the cause of the male infertility, we examined the cauda epididymis of *Ccdc38*^*+/+*^ and *Ccdc38*^*-/-*^ mice by Hematoxylin and Eosin (H&E) staining, and found there was fewer spermatozoa in the epididymal lumen of *Ccdc38*^*-/-*^ mice compared with *Ccdc38*^*+/+*^ mice (Fig. 3A). We then released the spermatozoa from epididymis, and found that the sperm number of *Ccdc38*^*-/-*^ mice was significantly less than that of *Ccdc38*^*+/+*^ mice (Fig. 3B), especially the motile spermatozoa decreased sharply (Fig. 3C). We further noticed *Ccdc38*^*-/-*^ spermatozoa bearing morphological aberrations, including abnormal nuclei and MMAF-like phenotype of short tail, curly tail, tailless (Fig. 3D). The ratio of spermatozoa with abnormal heads and flagella was shown in Fig. 3E. Scanning electron microscopy (SEM) detailed the morphological abnormalities of *Ccdc38*^*-/-*^ spermatozoa as follows (Fig. 3F): short tail (Type 1); disordered filaments (Type 2); impaired spermatozoa head (Type 4); curly tail (Type 5). Therefore, the knockout of *Ccdc38* results in MMAF-like phenotype in mice.

**Fig. 3.**
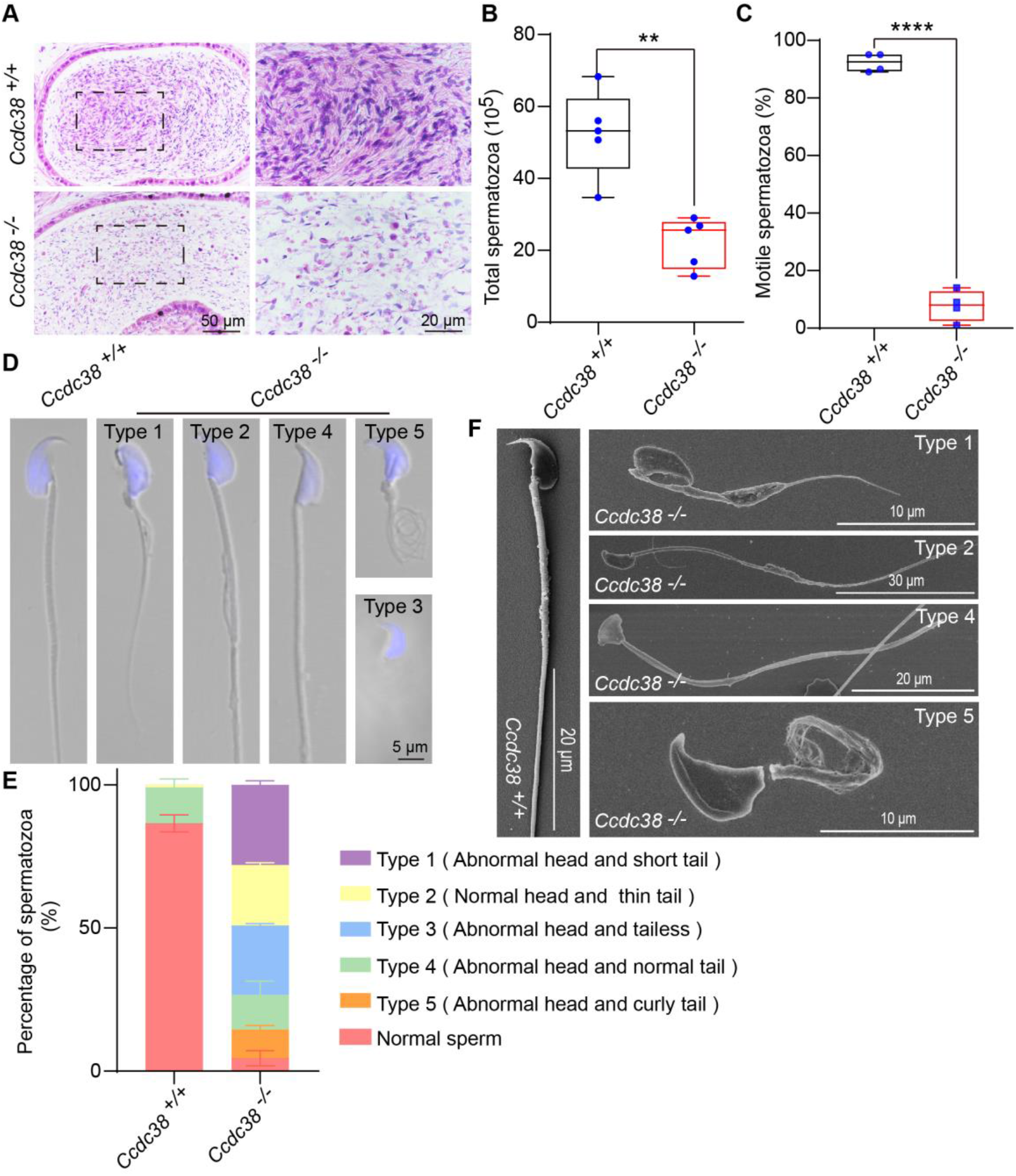
*Ccdc38* knockout results in MMAF. (A) H&E staining of the caudal epididymis. (B) The sperm number of *Ccdc38*^*+/+*^ and *Ccdc38*^*-/-*^ mice (n=5), ^**^P < 0.001. (C) The ratio of motile spermatozoa in *Ccdc38*^*+/+*^ and *Ccdc38*^*-/-*^ mice (n=5), ^****^P < 0.0001. (D) The single-sperm immunofluorescence analysis of *Ccdc38*^*+/+*^ and *Ccdc38*^*-/-*^ mice, nucleus was stained with DAPI. There were 5 phenotypes of the sperm: short tail, disordered tail, tailless, abnormal nuclei and curly tail. (E) The percentage of different spermatozoa in *Ccdc38*^*+/+*^ and *Ccdc38*^*-/-*^ caudal epididymis. (F) Scanning electron microscopy analysis of sperm from epididymis of *Ccdc38*^*+/+*^ and *Ccdc38*^*-/-*^ mice. It’s same as the immunofluorescence analysis except of tailless.

### Spermiogenesis is defected in *Ccdc38*^*-/-*^ mice

To further investigate why *Ccdc38* knockout leads to MMAF-like phenotype, we first used Periodic Acid Schiff (PAS) staining to determine at which stage the defect occurred. In *Ccdc38*^*+/+*^ mice testis section, round spermatid differentiated into elongating spermatids at stage IX, while there still were round spermatid and mature sperm at stage IX in *Ccdc38*^*-/-*^ mice testis (Fig. 4A). In order to delineate the detail defects of *Ccdc38*^*-/-*^ spermatids, we analyzed step 1-16 spermatids of both *Ccdc38*^*+/+*^ and *Ccdc38*^*-/-*^ mice, and found that in steps 1-8, the morphology of acrosome and nucleus of *Ccdc38*^*-/-*^ spermatids were similarly to that of the WT. In *Ccdc38*^*+/+*^ mice, spermatid head began elongation and mature from step 9, while in *Ccdc38*^*-/-*^ mice, spermatid head were abnormally elongated in step 9, eventually formed abnormal sperm at step16 (Fig. 4B). These results mean CCDC38 plays essential role during spermiogenesis.

**Fig. 4.**
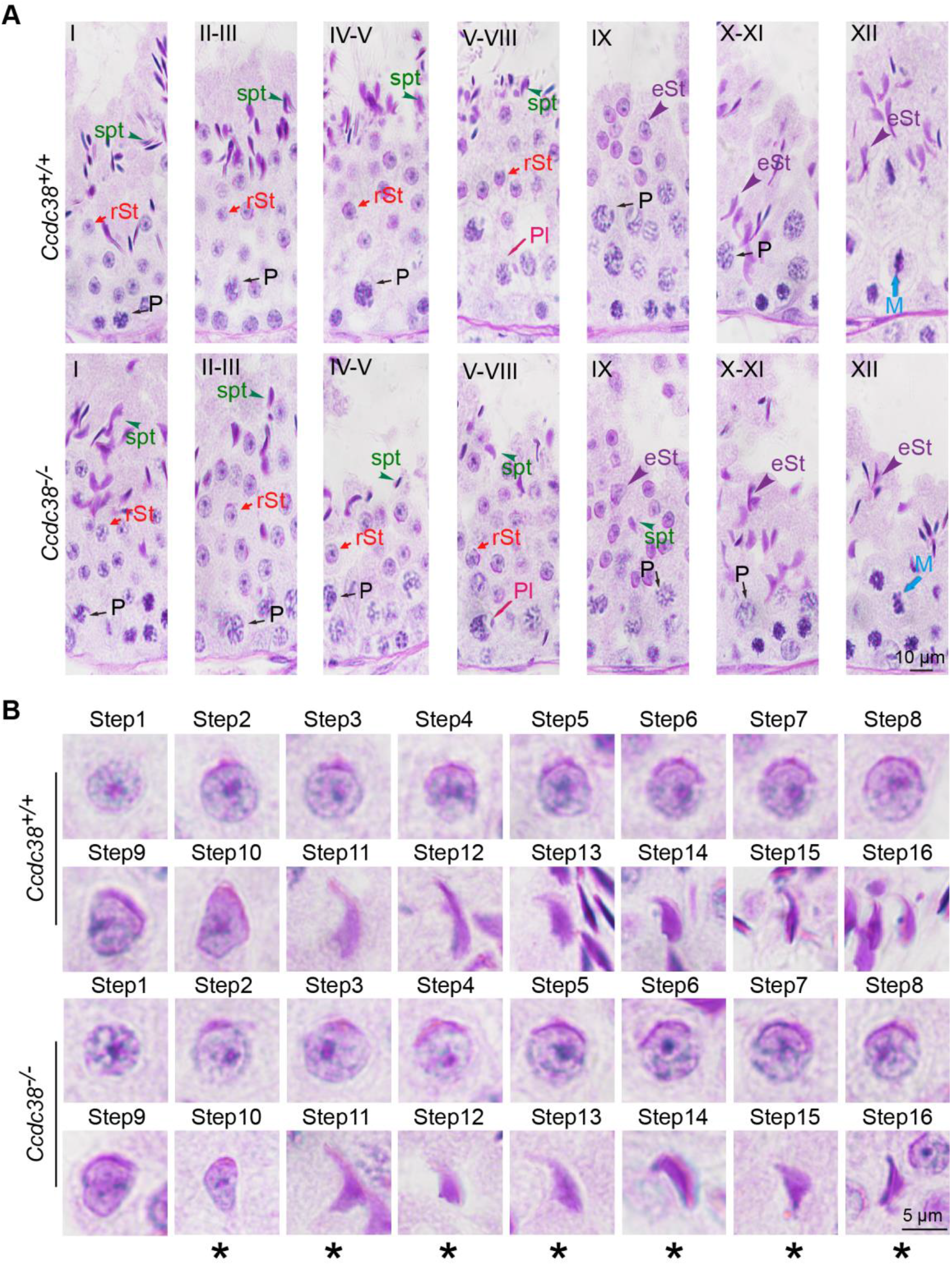
Spermiogenesis is defected in *Ccdc38*^*-/-*^ mice. (A) PAS staining of *Ccdc38*^*-/-*^ testis sections showed abnormal sperm nuclear shape. P: pachytene, rst: round spermatid, spt: spermatozoa, M: meiotic spermatocyte, In: spermatogonia. (B) PAS staining of spermatid at different steps from *Ccdc38*^*+/+*^ and *Ccdc38*^*-/-*^ mice. Asterisks indicated abnormal spermatid shape were found at step 10.

### Flagellum is disorganized and Manchette is ectopically placed in *Ccdc38*^*-/-*^ spermatids

To study the causes of abnormal sperm morphology after *Ccdc38* depletion, H&E staining was used to detect the morphology of seminiferous tubules between *Ccdc38*^*+/+*^ and *Ccdc38*^*-/-*^ mice. Compared with *Ccdc38*^*+/+*^ testis, obvious shortened tail and tailless sperm could be detected in *Ccdc38*^*-/-*^ testis (Fig. 5A). Immunofluorescence staining for acetylated TUBULIN, the specific flagellum marker, further confirmed the flagellum biogenesis defects in *Ccdc38*^*-/-*^ testis (Fig. 5B). We conducted immunofluorescence analysis of both PNA and α/β TUBULIN to determine which stages were affected by *Ccdc38* knockout, and found that the flagella of *Ccdc38*^*-/-*^ spermatids were shorter and curly from stage IV-V than that of *Ccdc38*^*+/+*^ spermatids (Fig. 5C). By using transmission electron microscopy (TEM), we observed that the Outer Dense Fibrous (ODF), Fibrous Sheath (FS) and mitochondria sheath were also abnormally organized in the *Ccdc38* KO elongating spermatids (Fig. 5D).

**Fig. 5.**
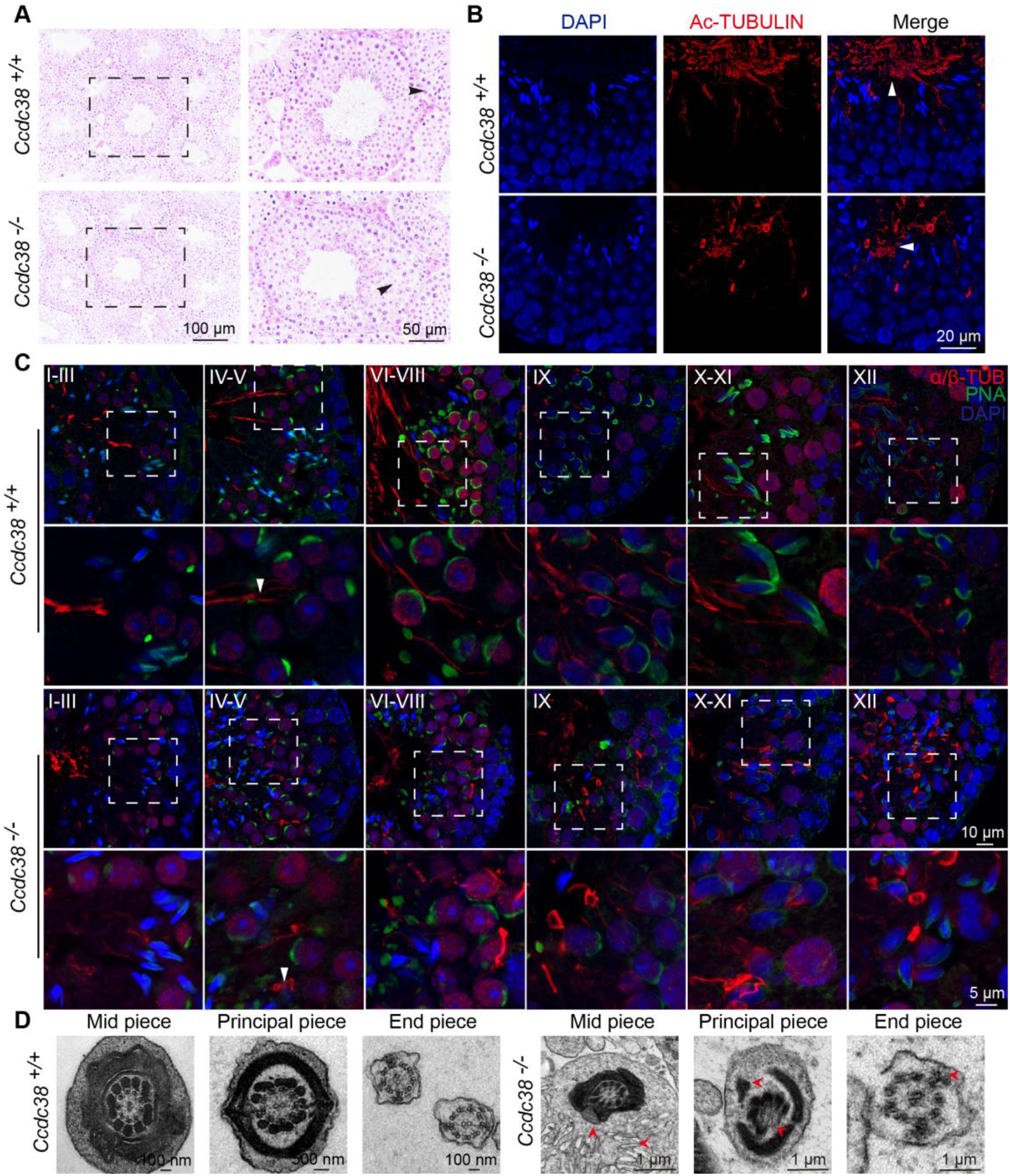
Flagellum is disorganized in *Ccdc38*^*-/-*^ spermatids. (A) The histology of the seminiferous tubules from *Ccdc38*^*+/+*^ and *Ccdc38*^*-/-*^ male mice. Arrows indicated the abnormal sperm. (B) Immunofluorescence analysis of AC-TUBULIN (red) antibodies from *Ccdc38*^*-/-*^ mice testes showed flagellar defects. Nucleus was stained with DAPI (blue), white arrows indicated the abnormal flagellum. (C) Immunofluorescence analysis of α/β-TUBULIN (red) and PNA lectin (green) to identify sperm flagellum biogenesis. White arrows indicated the short tail at stage IV-V compare with control group. (D) Cross sections of *Ccdc38*^*-/-*^ sperm tail to reveal the disorganization of axonemal microtubules and tail accessory structures (mitochondrial and fibrous sheath, outer dense fiber, red arrows indicated).

When spermatids were elongated, the sperm head was abnormal, indicating that the manchette might be abnormally formed (Fig. 5C). Manchette is important for sperm head shaping (Wei and Yang, 2018). So, we scrutinized manchette structure, and found the manchette of *Ccdc38*^*-/-*^ spermatids were roughly normal at steps 8-10, but from steps 11-12, they displayed abnormally longer than that of the control mice (Fig. 6A). We also used TEM to detect the manchette, *Ccdc38* knockout spermatids became abnormally elongated from step 11 but not in the control spermatids (Fig. 6B). In support of these result, we found that CCDC38 co-localized with α-TUBULIN at manchette in the control mice (Fig. 6C). All these results suggest that CCDC38 should be involved in flagellum biogenesis.

**Fig. 6.**
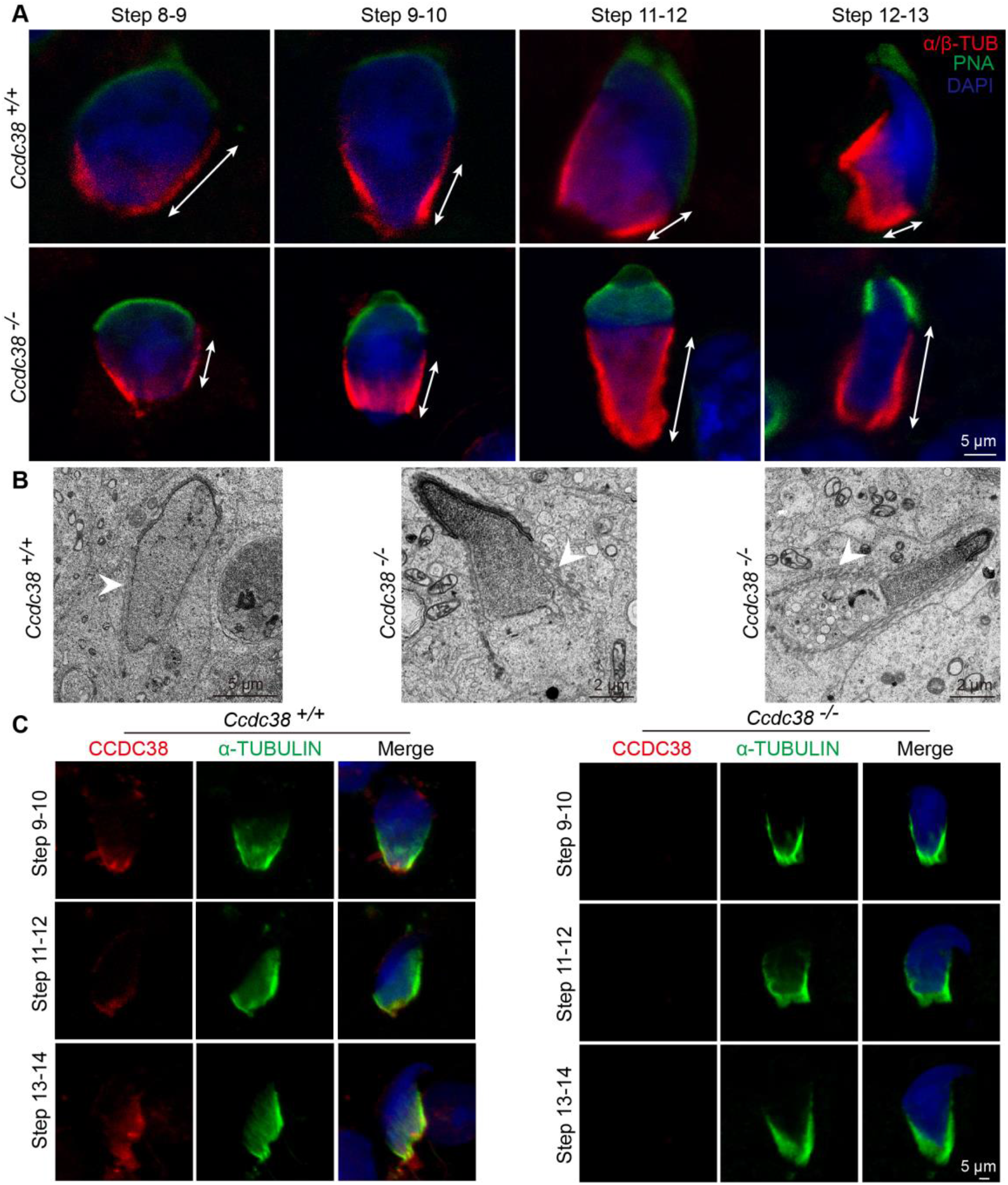
Manchette is ectopically placed in *Ccdc38*^*-/-*^ spermatids. (A) Abnormal manchette elongation in *Ccdc38*^*-/-*^ spermatids. Spermatids from different manchette containing steps were stained with anti α/β-TUBULIN antibody (red) and PNA lectin (green, acrosome marker) to visualize manchette. *Ccdc38*^*-/-*^ spermatids display abnormal elongation of the manchette. (B) TEM revealed that the manchette of the elongating spermatids (steps 9-11) of *Ccdc38*^*-/-*^ mice were ectopically placed (white arrows indicated). (C) Localization of CCDC38 in different stage germ cells. The immunofluorescence of CCDC38 and α-TUBULIN at developing germ cells. Manchette was stained with anti-α-TUBULIN antibody, nucleus was stained with DAPI.

### CCDC38 interacts with IFT88

It has been reported that CCDC42, IFT88 and KIF3A are involved in the anterograde transportation during flagellum biogenesis (Wu et al., 2021). To test whether CCDC38 also participates in anterograde transportation by interacting with IFT complexes, such as IFT88 and IFT20, we co-transfected pCSII-MYC-IFT88 or pRK-FLAG-IFT20 with pEGFP-C1-CCDC38 to the HEK293T cells, then immunoprecipitated CCDC38 with anti-GFP antibody, and found that IFT88 could be immunoprecipitated by CCDC38 (Fig. 7A), but not IFT20 (Fig. 7B). We also detected their expression level in *Ccdc38*^*+/+*^ and *Ccdc38*^*-/-*^ mice testis, and found IFT88 and IFT20 expression were all obviously decreased in *Ccdc38*^*-/-*^ mice testis (Fig. 7C, D). Then we detected the distribution of IFT88 in spermatids at different steps, and found that IFT88 was presented in the manchette and elongating sperm tails in *Ccdc38*^*+/+*^ mice, while in the *Ccdc38*^*-/-*^ spermatids, IFT88 still trapped close to the nucleus with a puncta-like structure (Fig. 7E). Therefore, CCDC38 might regulate sperm flagellum biogenesis by interacting with IFT B complexes.

**Fig. 7.**
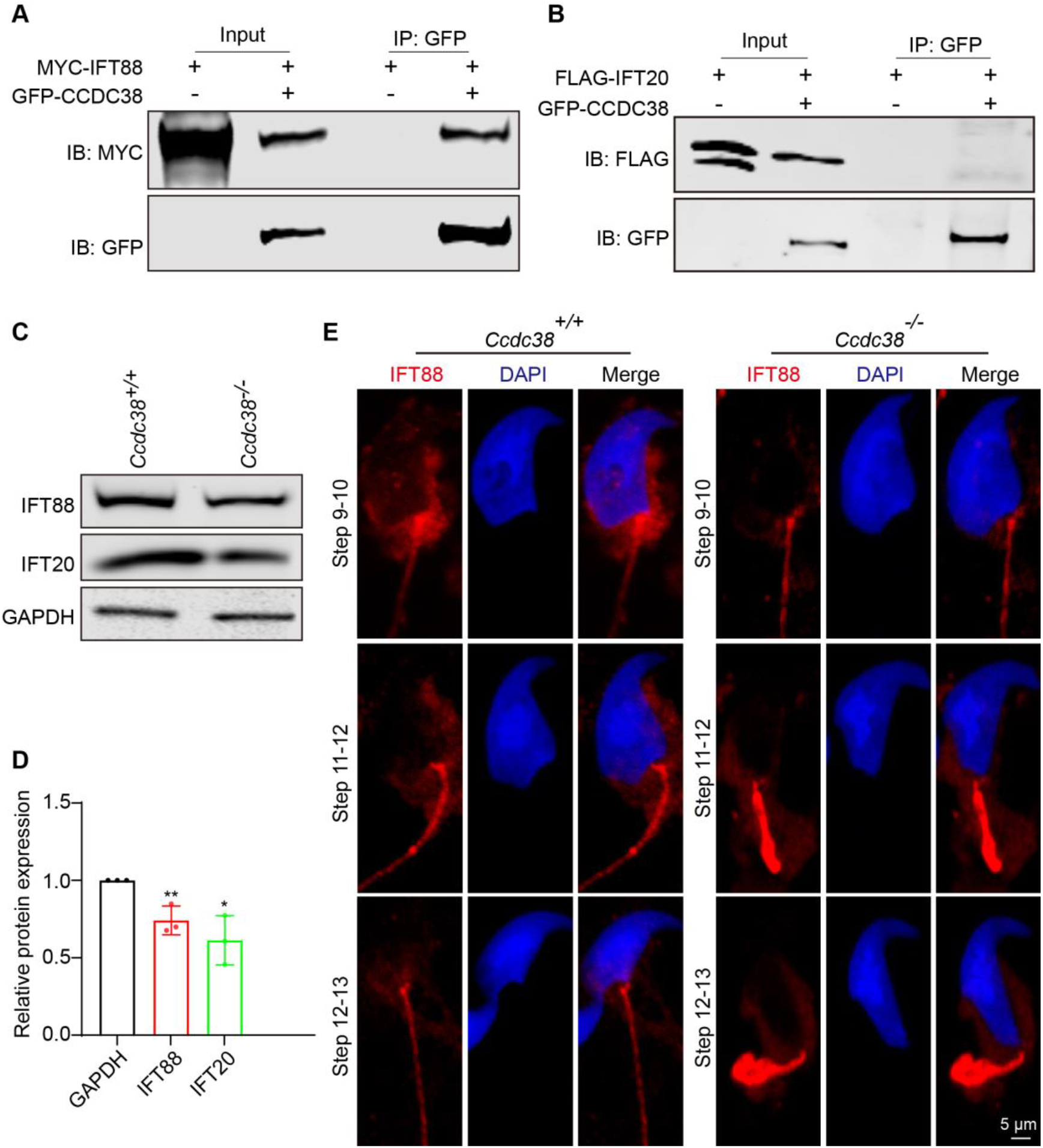
CCDC38 interacts with IFT88. (A) CCDC38 interacts with IFT88. pCSII-MYC-IFT88 were transfected into HEK293T cells with pEGFP-C1-CCDC38, forty-eight hours after transfection, cells were collected for immunoprecipitation with anti-GFP antibody, and analyzed with anti-GFP or anti-MYC antibodies, respectively. (B) CCDC38 cannot interact with IFT20. pRK-FLAG-IFT20 were transfected into HEK293T cells with pEGFP-C1-CCDC38, forty-eight hours after transfection, cells were collected for immunoprecipitation with anti-GFP antibody, and analyzed with anti-GFP or anti-FLAG antibodies, respectively. (C) Western blotting analysis to show IFT88, IFT20 protein levels in *Ccdc38*^*+/+*^ and *Ccdc38*^*-/-*^ mice testis lysates. GAPDH served as a loading control. (D) The quantitative results of western blotting. ^**^P < 0.001, ^*^P < 0.01 indicates a significant difference (t-test). (E) Immunofluorescence of IFT88 (red) and DAPI (blue) in spermatids at different stages from *Ccdc38*^*+/+*^ and *Ccdc38*^*-/-*^ mice.

### ODF transportation is defected in *Ccdc*38 knockout spermatids

It has been reported that ODF1 and ODF2 could interact with CCDC42, and they are found to be involved in the formation of male germ cell cytoskeleton (Tapia Contreras and Hoyer-Fender, 2019). To study the relationship between CCDC38 and ODF2, reciprocal coimmunoprecipitation assays were carried out. we transfected pCDNA-HA-ODF2 plasmid and pEGFP-C1-CCDC38 plasmid in to HEK293T cells, CCDC38 and ODF2 were able to interact with the other in reciprocal immunoprecipitation experiments (Fig. 8A), suggesting CCDC38 might interact with ODF2.

**Fig. 8.**
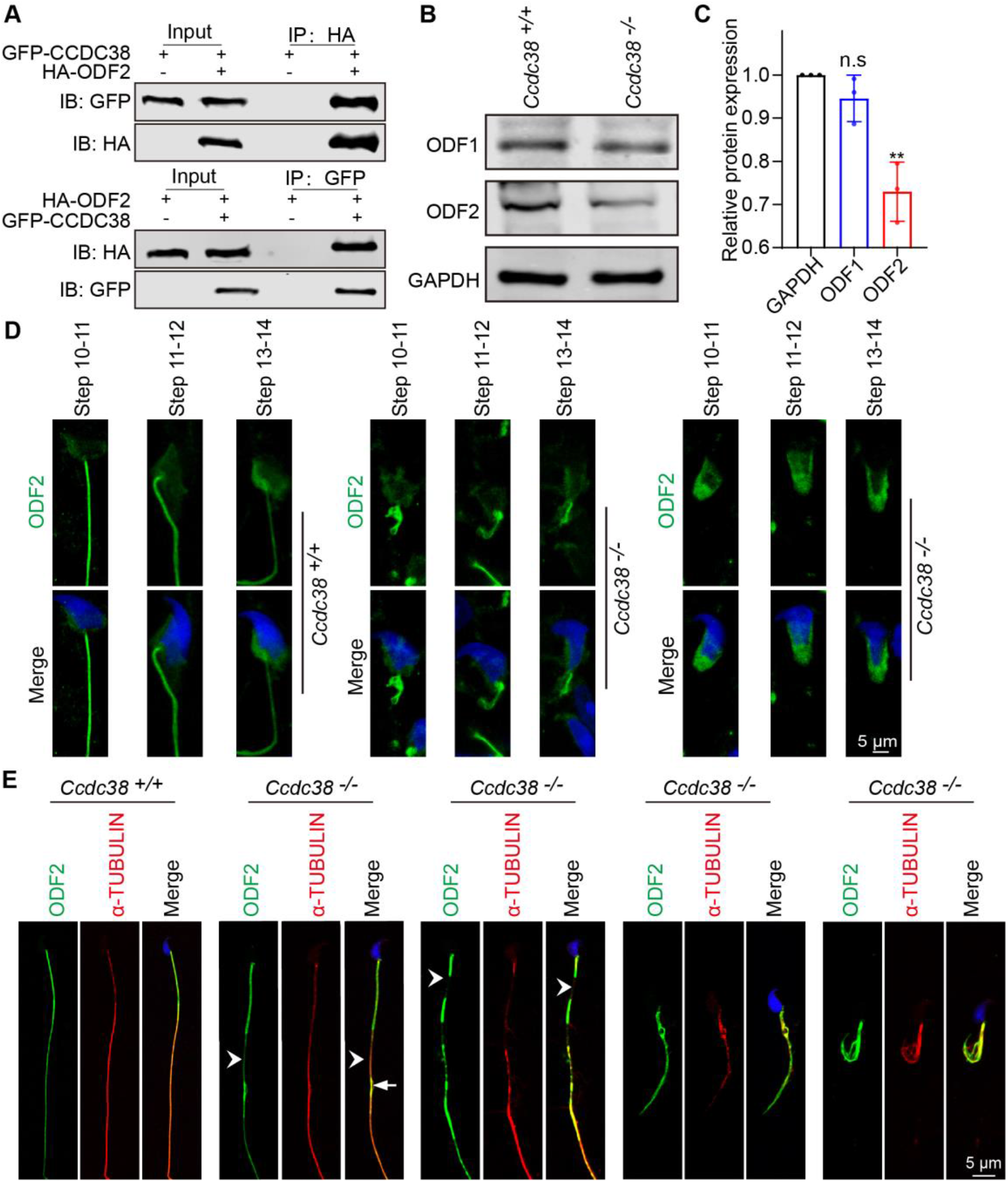
ODF transportation is defected in *Ccdc*38 knockout spermatids. (A) CCDC38 interacted with ODF2. pCDNA-HA-ODF2 and pEGFP-C1-CCDC38 were transfected into HEK293T cells, forty-eight hours after transfection, cells were collected for immunoprecipitation with anti-GFP or anti-HA antibodies, and then analyzed with anti-GFP or anti-HA antibodies, respectively. (B) Western blotting analysis to show ODF1, ODF2 protein levels in *Ccdc38*^*+/+*^ and *Ccdc38*^*-/-*^ mice testis lysates. GAPDH served as a loading control. ODF2 protein level was decreased. (C) The quantitative results of western blot. ^**^P < 0.01 indicates a significant difference (t-test). (D) The localization of ODF2 in testis germ cells. Testicular germ cells were stained with anti-ODF2 antibody (green), ODF2 was localized in spermatid flagellum and manchette in *Ccdc38*^*-/-*^ or *Ccdc38*^*+/+*^ germ cells. (E) Immunofluorescence of ODF2 (green) and α-TUBULIN (red) in spermatids from *Ccdc38*^*+/+*^ and *Ccdc38*^*-/-*^ mice. Nucleus was stained with DAPI (blue), white arrows indicated the discontinuous, punctiform short, white arrowhead indicated the tenuous axoneme.

As the main cytoskeleton protein in the ODFs, ODF2 is essential for sperm flagellum integrity and beating (Donkor et al., 2004, Ito et al., 2019, Fawcett, 1975). We examined the effect of *Ccdc38* knockout on ODF1 and ODF2 protein levels, and found that ODF2, but not ODF1, was significantly decreased in *Ccdc38*^*-/-*^ testicular extracts (Fig. 8B, C). Then, we used immunofluorescence to detect the expression of ODF2 in spermatids and epididymal spermatozoa. We found that ODF2 localized on manchette along with the sperm tail in elongated spermatids of *Ccdc38*^*+/+*^ mice, whereas ODF2 was detected on manchette without tail staining in most of elongated spermatids (Fig. 8D). Of note, ODF2 co-localized with α-TUBULIN on the midpiece and principal piece of *Ccdc38*^*+/+*^ sperm tail, while ODF2 signal displayed discontinuous, punctiform short or curly on *Ccdc38* knockout spermatozoa (Fig. 8E), suggesting that the defects of ODFs in *Ccdc38* knockout spermatozoa might come from a defect of ODF2 transportation during spermiogenesis.

## Discussion

*Ccdc38* is a testis specific expression gene (Lin et al., 2016), but its role during spermiogenesis has not been investigated yet. In order to study its role during spermiogenesis, we generated *Ccdc38*^*-/-*^ mouse model. *Ccdc38*^*-/-*^ male mice was sterile (Fig. 2F) due to significantly reduced spermatozoa number and motility (Fig. 3B, C), albeit with no significant size difference between *Ccdc38*^*+/+*^ and *Ccdc38*^*-/-*^ testes (Fig. 2G).

The manchette is a transient structure in developing germ cells, which is required for sperm nuclear condensation and flagellum biogenesis (Wei and Yang, 2018). It provides the structural basis for intra-manchette transport (IMT), IMT transfers structural and functional proteins to the basal body and is essential for nucleo-cytoplasmic transport (Kierszenbaum, 2002, Kierszenbaum et al., 2002). As an IMT component, CCDC42 localizes to the manchette, connecting piece and sperm tail during spermiogenesis, and it can interact with ODF1, ODF2 to regulate germ cell cytoskeleton formation (Tapia Contreras and Hoyer-Fender, 2019, Pasek et al., 2016). Here, we found that CCDC38 could interact with CCDC42, and co-localized with CCDC42 on centrosome in Hela cells (Fig. 1A, B, C). In addition, CCDC38 were found to localize on manchette and sperm tail (Fig. 1D), and it interacted with ODF2 (Fig. 8A). ODF2 is a component of outer dense fibers, and it is important for sperm flagellum assembly, the knockout of this gene leads to preimplantation lethal, and even the absence of a single copy of this gene results in sperm neck-midpiece separation (Qian et al., 2016, Tarnasky et al., 2010). Here, we found that once *Ccdc38* was knocked out, the protein level of ODF2 was decreased in testis (Fig. 8B, C) and its distribution was disturbed in flagella (Fig. 8D, E). Thus, CCDC38 either works as a partner of ODF2 to keep its stability or participates in IMT to intermediate ODF2 transportation during flagellum biogenesis. Since CCDC38 also interacted with CCDC42, we prefer the second one, and this possibility is also supported by its interaction with IFT88.

In addition to IMT, the intra-flagellar transport (IFT) is also required for flagellum biogenesis. IFT is responsible for sperm-protein transportation during the development of the flagella. During IFT, cargoes are transported from the basal body to the tips of the flagellum and then back to the sperm head along the axoneme (Scholey, 2003, Taschner and Lorentzen, 2016, Ishikawa and Marshall, 2017). IFT88 is an IFT B components, it presents in the heads and tails only in step 15, and no longer being detected in mature sperm (San Agustin et al., 2015), it can interact with kinesin to regulate the anterograde transport along axoneme (Rosenbaum and Witman, 2002). Worked as an IFT88 interacting protein (Fig. 7A), CCDC38 may also participate in the anterograde transport along the flagellum. Thus, CCDC38 may interact with both CCDC42 and IFT88 to regulate cargoes transportation by IMT and IFT during flagellum biogenesis.

In summary, we identified a new CCDC42 interacting protein, CCDC38, which is essential for spermiogenesis and flagellum biogenesis, the knockout of this gene results in MMAF-like phenotype in mice. Since these genes are evolutionary conserved in human beings, we believe that some mutations of these genes should be existed in MMAF patients, albeit we do not find them right now.

## Materials and methods

### Animals

The mice *Ccdc38* gene is 1692 bp and contains 16 exons. The knockout mice of *Ccdc38* were generated by CRISPER-Cas9 system from Cyagen Biosciences. The genotyping primers for knockout were as follows: F1: GTAGCTGTTTCTAAGCGATCATCA, R1: ACTAGGTACCTCAAGCTGGTTTAGA, and for WT mice, the specific primers were: F1: GTAGCTGTTTCTAAGCGATCATCA, R2: GTCATGGGACAGATGTGGAACTA.

All the animal experiments were performed according to approved institutional animal care and use committee (IACUC) protocols (# 08-133) of the Institute of Zoology, Chinese Academy of Sciences.

### Antibodies

Mouse anti-GFP antibody (1:1000, M20004L, Abmart), rabbit anti-MYC antibody (1:1000, BE2011, Abmart), ODF2 antibody (12058-1-AP, Proteintech) was used at a dilution at 1:1000 for western blotting and 1: 200 for immunofluorescence. Mouse anti-α-TUBULIN antibody (1:200, AC012, Abclonal) for immunofluorescence. Mouse anti-GAPDH antibody (1:10000, AC002, Abclonal) for western blotting. Mouse anti-ODF1 antibody (1:500, sc-390152, santa) for western blotting. Mouse anti-CCDC38 were generated from Dia-an Biotech (Wuhan, China). The Alexa Fluor 488 conjugate of lectin PNA (1:400, L21409, Thermo Fisher), the Mito-Tracker Deep Red 633 (1:1000, M22426, Thermo Fisher) were used for immunofluorescence. The secondary antibodies were goat anti rabbit FITC (1:200, ZF-0311, Zhong Shan Jin Qiao), goat anti TRITC (1:200, ZF-0316, Zhong Shan Jin Qiao), goat anti mouse FITC (1:200, ZF-0312, Zhong Shan Jin Qiao), goat anti rabbit TRITC (1:200, ZF0313, Zhong Shan Jin Qiao).

### Immunoblotting

As previously reported (Liu et al., 2016), testis albuginea was peeled and added in RIPA buffer supplemented with 1mM phenyl methyl sulfonyl fluoride (PMSF) and PIC (Roche Diagnostics, 04693132001), the solution was sonicated transiently and then on the ice for 30 min. The samples were centrifuged at 12000 rpm for 15 min at 4°C. Then, the supernatant was collected at a new tube. The protein lysates were electrophoresed and electrotransfered, then incubated with primary antibody and second antibody, next the membrane was scanned via an Odyssey infrared imager (LI-COR Biosciences, Lincoln, NE, RRID:SCR_014579).

### Immunoprecipitation

Transfected cells were lysed in a lysis buffer (50mM HEPES, PH 7.4, 250mM NaCl, 0.1% NP-40 containing PIC and PMSF) on ice for 30 min, and centrifugated at 12000 rpm at 4°C for 15 min, cell lysates were incubated with primary antibody overnight at 4 °C, next incubated with protein A for 2h at 4°C, then washed 3 times with lysed buffer and subjected to immunoblotting analysis.

### Epididymal sperm count

The cauda epididymis was isolated from 8 weeks mice. Sperm was released from the cauda epididymis with HTF and incubated at 37°C for 15 min. Then the medium was diluted at 1:100 and counted the sperm number with hemocytometer.

### Tissue collection and histological analysis

As previously reported (Wang et al., 2018), the testes were dissected after euthanasia, and fixed with Bouin’s fixative for 24h at 4 °C, then the testes were dehydrated with graded ethanol and embedded in paraffin. The 5um sections were cutted and covered on glass slides. Sections were stained with H&E and PAS for histological analysis after deparaffinization.

### Transmission electron microscopy

The methods were as reported previously with some modifications (Liu et al., 2016). The testis from WT and *Ccdc38* depletion mice testis and epididymis were dissected and fixed in 2.5% glutaraldehyde in 0.1 M cacodylate buffer at 4°C overnight. After washing in 0.1 M cacodylate buffer, samples were cutted into small pieces, then immersed in 1% OsO4 for 1h at 4°C. Samples were dehydrated through a graded acetone series and embedded in resin for staining. Ultrathin sections were cutted and stained with uranyl acetate and lead citrate, images were acquired and analyzed using a JEM-1400 transmission electron microscope.

### Scanning electron microscopy

The sperm were released from epididymis in HTF at 37°C 15 min, centrifugated 5 min at 500 g, then washed twice with PB, and fixed in 2.5% glutaraldehyde solution overnight, and dehydrated in a graded ethanol, subjected to drying and coated with gold. The images were acquired and analyzed using SU8010 scanning electron microscope.

### Immunofluorescence

The testis albuginea was peeled and incubated with collagenase IV and hyaluronidase in PBS for 15 min at 37°C, then washed twice with PBS. Next, fixed with 4% PFA 5 min, and then coated on slide glass to dry out. The slides were washed with PBS three times and then treated with 0.5% TritonX-100 for 5 min, and blocked with 5% BSA for 30 min. Added the primary antibodies and incubated at 4°C overnight, followed by incubating with second antibody and DAPI. The images were taken using a LSM880 and Sp8 microscopes.

### Statistical Analysis

All data are presented as the mean ±SEM. The statistical significance of the differences between the mean values for the various genotypes was measured by Student’s t-tests with paired, 2-tailed distribution. The data were considered significant when the P-value was less than 0.05(^*^), 0.01(^**^) or 0.001(^***^).

## Acknowledgements

This work was funded by the National Natural Science Foundation of China (grants 91649202), the Strategic Priority Research Program of the Chinese Academy of Sciences (grant XDA16020701), and the National Key R&D Program of China (grant 2016YFA0500901).

## Authors Contributions

RDZ and BBW performed most of the experiments and wrote the manuscript. CL, XGW, LYW, XS and YHC performed part of the experiment. WL supervised the whole project and revised the manuscript.

## Compliance with ethical standards

All animal experiments were performed according to approved institutional animal care and use committee (IACUC) protocols (#08-133) of the Institute of Zoology, Chinese Academy of Sciences. All surgery was performed under sodium pentobarbital anesthesia, and every effort was made to minimize suffering.

## References

Azizi, F. & Ghafouri-Fard, S. 2017. Outer Dense Fiber Proteins: Bridging between Male Infertility and Cancer. Arch Iran Med, 20, 320–325.

Coutton, C., Escoffier, J., Martinez, G., Arnoult, C. & Ray, P. F. 2015. Teratozoospermia: spotlight on the main genetic actors in the human. Hum Reprod Update, 21, 455–85.

Donkor, F. F., Monnich, M., Czirr, E., Hollemann, T. & Hoyer-Fender, S. 2004. Outer dense fibre protein 2 (ODF2) is a self-interacting centrosomal protein with affinity for microtubules. J Cell Sci, 117, 4643–51.

Fawcett, D. W. 1975. The mammalian spermatozoon. Dev Biol, 44, 394–436.

Firat-Karalar, E. N., Sante, J., Elliott, S. & Stearns, T. 2014. Proteomic analysis of mammalian sperm cells identifies new components of the centrosome. J Cell Sci, 127, 4128–33.

Freitas, M. J., Vijayaraghavan, S. & Fardilha, M. 2017. Signaling mechanisms in mammalian sperm motility. Biol Reprod, 96, 2–12.

Inaba, K. 2011. Sperm flagella: comparative and phylogenetic perspectives of protein components. Mol Hum Reprod, 17, 524–38.

Ishikawa, H. & Marshall, W. F. 2017. Intraflagellar Transport and Ciliary Dynamics. Cold Spring Harb Perspect Biol, 9.

Ito, C., Akutsu, H., Yao, R., Yoshida, K., Yamatoya, K., Mutoh, T., Makino, T., Aoyama, K., Ishikawa, H., Kunimoto, K., Tsukita, S., Noda, T., Kikkawa, M. & Toshimori, K. 2019. Odf2 haploinsufficiency causes a new type of decapitated and decaudated spermatozoa, Odf2-DDS, in mice. Sci Rep, 9, 14249.

Jiao, S. Y., Yang, Y. H. & Chen, S. R. 2021. Molecular genetics of infertility: loss-of-function mutations in humans and corresponding knockout/mutated mice. Hum Reprod Update, 27, 154–189.

Kierszenbaum, A. L. 2002. Intramanchette transport (IMT): managing the making of the spermatid head, centrosome, and tail. Mol Reprod Dev, 63, 1–4.

Kierszenbaum, A. L., Gil, M., Rivkin, E. & Tres, L. L. 2002. Ran, a GTP-binding protein involved in nucleocytoplasmic transport and microtubule nucleation, relocates from the manchette to the centrosome region during rat spermiogenesis. Mol Reprod Dev, 63, 131–40.

Kim, Y. H., Mcfarlane, J. R., O’bryan, M. K., Almahbobi, G., Temple-Smith, P. D. & De Kretser, D. M. 1999. Isolation and characterization of rat sperm tail outer dense fibres and comparison with rabbit and human spermatozoa using a polyclonal antiserum. J Reprod Fertil, 116, 345–53.

Lehti, M. S. & Sironen, A. 2017. Formation and function of sperm tail structures in association with sperm motility defects. Biol Reprod, 97, 522–536.

Li, L., Feng, F., Wang, Y., Guo, J. & Yue, W. 2020. Mutational effect of human CFAP43 splice-site variant causing multiple morphological abnormalities of the sperm flagella. Andrologia, 52, e13575.

Li, Y., Sha, Y., Wang, X., Ding, L., Liu, W., Ji, Z., Mei, L., Huang, X., Lin, S., Kong, S., Lu, J., Qin, W., Zhang, X., Zhuang, J., Tang, Y. & Lu, Z. 2019. DNAH2 is a novel candidate gene associated with multiple morphological abnormalities of the sperm flagella. Clin Genet, 95, 590–600.

Lin, S. R., Li, Y. C., Luo, M. L., Guo, H., Wang, T. T., Chen, J. B., Ma, Q., Gu, Y. L., Jiang, Z. M. & Gui, Y. T. 2016. Identification and characteristics of the testes-specific gene, Ccdc38, in mice. Mol Med Rep, 14, 1290–6.

Liu, C., Miyata, H., Gao, Y., Sha, Y., Tang, S., Xu, Z., Whitfield, M., Patrat, C., Wu, H., Dulioust, E., Tian, S., Shimada, K., Cong, J., Noda, T., Li, H., Morohoshi, A., Cazin, C., Kherraf, Z. E., Arnoult, C., Jin, L., He, X., Ray, P. F., Cao, Y., Toure, A., Zhang, F. & Ikawa, M. 2020. Bi-allelic DNAH8 Variants Lead to Multiple Morphological Abnormalities of the Sperm Flagella and Primary Male Infertility. Am J Hum Genet, 107, 330–341.

Liu, C., Wang, H., Shang, Y., Liu, W., Song, Z., Zhao, H., Wang, L., Jia, P., Gao, F., Xu, Z., Yang, L., Gao, F. & Li, W. 2016. Autophagy is required for ectoplasmic specialization assembly in sertoli cells. Autophagy, 12, 814–32.

Liu, W., He, X., Yang, S., Zouari, R., Wang, J., Wu, H., Kherraf, Z. E., Liu, C., Coutton, C., Zhao, R., Tang, D., Tang, S., Lv, M., Fang, Y., Li, W., Li, H., Zhao, J., Wang, X., Zhao, S., Zhang, J., Arnoult, C., Jin, L., Zhang, Z., Ray, P. F., Cao, Y. & Zhang, F. 2019. Bi-allelic Mutations in TTC21A Induce Asthenoteratospermia in Humans and Mice. Am J Hum Genet, 104, 738–748.

Pasek, R. C., Malarkey, E., Berbari, N. F., Sharma, N., Kesterson, R. A., Tres, L. L., Kierszenbaum, A. L. & Yoder, B. K. 2016. Coiled-coil domain containing 42 (Ccdc42) is necessary for proper sperm development and male fertility in the mouse. Dev Biol, 412, 208–18.

Pereira, R., Sa, R., Barros, A. & Sousa, M. 2017. Major regulatory mechanisms involved in sperm motility. Asian J Androl, 19, 5–14.

Perles, Z., Cinnamon, Y., Ta-Shma, A., Shaag, A., Einbinder, T., Rein, A. J. & Elpeleg, O. 2012. A human laterality disorder associated with recessive CCDC11 mutation. J Med Genet, 49, 386–90.

Priyanka, P. P. & Yenugu, S. 2021. Coiled-Coil Domain-Containing (CCDC) Proteins: Functional Roles in General and Male Reproductive Physiology. Reprod Sci.

Qian, X., Wang, L., Zheng, B., Shi, Z. M., Ge, X., Jiang, C. F., Qian, Y. C., Li, D. M., Li, W., Liu, X., Yin, Y., Zheng, J. T., Shen, H., Wang, M., Guo, X. J., He, J., Lin, M., Liu, L. Z., Sha, J. H. & Jiang, B. H. 2016. Deficiency of Mkrn2 causes abnormal spermiogenesis and spermiation, and impairs male fertility. Sci Rep, 6, 39318.

Rosenbaum, J. L. & Witman, G. B. 2002. Intraflagellar transport. Nat Rev Mol Cell Biol, 3, 813–25.

San Agustin, J. T., Pazour, G. J. & Witman, G. B. 2015. Intraflagellar transport is essential for mammalian spermiogenesis but is absent in mature sperm. Mol Biol Cell, 26, 4358–72.

Scholey, J. M. 2003. Intraflagellar transport. Annu Rev Cell Dev Biol, 19, 423–43.

Sha, Y.-W., Xu, X., Mei, L.-B., Li, P., Su, Z.-Y., He, X.-Q. & Li, L. 2017. A homozygous CEP135 mutation is associated with multiple morphological abnormalities of the sperm flagella (MMAF). Gene, 633, 48–53.

Sha, Y., Xu, Y., Wei, X., Liu, W., Mei, L., Lin, S., Ji, Z., Wang, X., Su, Z., Qiu, P., Chen, J. & Wang, X. 2019. CCDC9 is identified as a novel candidate gene of severe asthenozoospermia. Syst Biol Reprod Med, 65, 465–473.

Sha, Y. W., Ding, L. & Li, P. 2014. Management of primary ciliary dyskinesia/Kartagener’s syndrome in infertile male patients and current progress in defining the underlying genetic mechanism. Asian J Androl, 16, 101–6.

Shen, Y., Zhang, F., Li, F., Jiang, X., Yang, Y., Li, X., Li, W., Wang, X., Cheng, J., Liu, M., Zhang, X., Yuan, G., Pei, X., Cai, K., Hu, F., Sun, J., Yan, L., Tang, L., Jiang, C., Tu, W., Xu, J., Wu, H., Kong, W., Li, S., Wang, K., Sheng, K., Zhao, X., Yue, H., Yang, X. & Xu, W. 2019. Loss-of-function mutations in QRICH2 cause male infertility with multiple morphological abnormalities of the sperm flagella. Nat Commun, 10, 433.

Silva, E., Betleja, E., John, E., Spear, P., Moresco, J. J., Zhang, S., Yates, J. R., 3RD, Mitchell, B.J. & Mahjoub, M. R. 2016. Ccdc11 is a novel centriolar satellite protein essential for ciliogenesis and establishment of left-right asymmetry. Mol Biol Cell, 27, 48–63.

Sironen, A., Shoemark, A., Patel, M., Loebinger, M. R. & Mitchison, H. M. 2020. Sperm defects in primary ciliary dyskinesia and related causes of male infertility. Cell Mol Life Sci, 77, 2029–2048.

Tang, S., Wang, X., Li, W., Yang, X., Li, Z., Liu, W., Li, C., Zhu, Z., Wang, L., Wang, J., Zhang, L., Sun, X., Zhi, E., Wang, H., Li, H., Jin, L., Luo, Y., Wang, J., Yang, S. & Zhang, F. 2017. Biallelic Mutations in CFAP43 and CFAP44 Cause Male Infertility with Multiple Morphological Abnormalities of the Sperm Flagella. Am J Hum Genet, 100, 854–864.

Tapia Contreras, C. & Hoyer-Fender, S. 2019. CCDC42 Localizes to Manchette, HTCA and Tail and Interacts With ODF1 and ODF2 in the Formation of the Male Germ Cell Cytoskeleton. Front Cell Dev Biol, 7, 151.

Tarnasky, H., Cheng, M., Ou, Y., Thundathil, J. C., Oko, R. & Van Der Hoorn, F. A. 2010. Gene trap mutation of murine outer dense fiber protein-2 gene can result in sperm tail abnormalities in mice with high percentage chimaerism. BMC Dev Biol, 10, 67.

Taschner, M. & Lorentzen, E. 2016. The Intraflagellar Transport Machinery. Cold Spring Harb Perspect Biol, 8.

Wang, L., Tu, Z., Liu, C., Liu, H., Kaldis, P., Chen, Z. & Li, W. 2018. Dual roles of TRF1 in tethering telomeres to the nuclear envelope and protecting them from fusion during meiosis. Cell Death Differ, 25, 1174–1188.

Wei, Y. L. & Yang, W. X. 2018. The acroframosome-acroplaxome-manchette axis may function in sperm head shaping and male fertility. Gene, 660, 28–40.

Wu, B., Yu, X., Liu, C., Wang, L., Huang, T., Lu, G., Chen, Z. J., Li, W. & Liu, H. 2021. Essential Role of CFAP53 in Sperm Flagellum Biogenesis. Front Cell Dev Biol, 9, 676910.

Yamaguchi, A., Kaneko, T., Inai, T. & Iida, H. 2014. Molecular cloning and subcellular localization of Tektin2-binding protein 1 (Ccdc 172) in rat spermatozoa. J Histochem Cytochem, 62, 286–97.

Young, S. A., Miyata, H., Satouh, Y., Kato, H., Nozawa, K., Isotani, A., Aitken, R. J., Baker, M. A. & Ikawa, M. 2015. CRISPR/Cas9-Mediated Rapid Generation of Multiple Mouse Lines Identified Ccdc63 as Essential for Spermiogenesis. Int J Mol Sci, 16, 24732–50.

